# Corticothalamic Layer 6 Controls Cortical Activity and Thalamic Firing Mode in a Bidirectional Manner

**DOI:** 10.1101/2025.07.17.665299

**Authors:** Ross Folkard, Emilio Ulises Isaías-Camacho, Alexander Groh

## Abstract

Corticothalamic layer 6 modulates information flow between cortical and thalamic circuits. Previous research reported contrasting inhibitory or excitatory effects of corticothalamic layer 6 on cortical dynamics, potentially reflecting technological discrepancies or physiological differences in corticothalamic layer 6 function. To resolve these discrepancies, we combined translaminar, multi-channel *in vivo* electrophysiology in the primary somatosensory cortex of the anaesthetised mouse with optogenetic stimulation across a range of stimulation regimes to manipulate firing rate and frequency of corticothalamic layer 6. Increasing corticothalamic layer 6 firing rates exerted a transition from inhibition to excitation across cortical layers. Furthermore, corticothalamic layer 6 activity imparted population synchrony onto distinct cortical subpopulations, independent of changes in overall corticothalamic layer 6 activity. In the thalamus, corticothalamic layer 6 modulated thalamic bursting in a bidirectional manner, dependent on optogenetic stimulation frequency. These results demonstrate that corticothalamic layer 6 in primary somatosensory cortex can bidirectionally modulate both cortical firing and thalamic firing mode, elucidating a more nuanced function of somatosensory corticothalamic layer 6 in thalamic and cortical signalling than previously recognised.

## Introduction

Corticothalamic layer 6 (L6CT) neurons play a key role in regulating activity in both the cortex and thalamus, exerting monosynaptic excitation and disynaptic inhibition onto their targets (Lam and Sherman 2010; Jurgens et al. 2012; Olsen et al. 2012; Bortone et al. 2014; Kim et al. 2014; Mease et al. 2014; Frandolig et al. 2019; Whilden et al. 2021). L6CT is known to bidirectionally modulate thalamic excitability depending on stimulation frequency (Crandall et al. 2015). However, whether L6CT similarly exerts activity-dependent control over cortical dynamics remains an open question.

In visual and somatosensory cortex *in vitro* experiments, L6CT has been shown to strongly excite layer 5 pyramidal neurons while weakly exciting layer 4 excitatory neurons (Kim et al., 2014). Additionally, L6CT provides feedforward inhibition to layers 4 and 5 via distinct GABAergic subtypes and can entrain cortical layers into rhythmic firing patterns (Kim et al., 2014; Frandolig et al., 2019; Olsen et al., 2012; Bortone et al., 2014; Adesnik, 2018). Despite these insights, *in vivo* studies using optogenetic activation of L6CT have reported conflicting findings, with some observing net excitatory effects and others reporting net inhibition, depending on the sensory modality and experimental conditions (Olsen et al. 2012; Guo et al. 2017; Ziegler et al. 2023; Dimwamwa et al. 2024).

This variability raises the question of whether these differences reflect genuine physiological distinctions between sensory cortices or are instead due to differences in experimental methodology. The few direct comparisons across modalities suggest that L6CT circuitry is largely conserved (Kim et al., 2014), implying that methodological factors may account for much of the observed heterogeneity. If this is the case, systematically varying L6CT stimulation parameters within a single cortical area should reproduce the range of effects seen across prior studies (Olsen et al. 2012; Guo et al. 2017; Ziegler et al. 2023; Dimwamwa et al. 2024).

In this present study, we examined how different L6CT stimulation regimes influence neuronal spiking in the primary somatosensory cortex (S1) and in ventroposterior lateral thalamus (VPL) in the hindlimb system of anaesthetised mice, with the central question of whether L6CT stimulation has a net excitatory, inhibitory, or bidirectional effect on ongoing spiking activity. We found that L6CT activity bidirectionally modulates layers 4 and 5, with low L6CT activity inducing inhibition and high L6CT activity driving excitation. Additionally, L6CT stimulation modulated thalamic firing mode in a frequency-dependent manner and thalamic firing rate as a function of L6CT activity. These findings provide new insights into how L6CT dynamically regulates cortical and thalamic activity, refining our understanding of its role in sensory processing.

## Materials and Methods

### Ethics Statement

The local governing body approved all of the following experimental procedures (Regierungspräsidium Karlsruhe, Germany, approval numbers: 35-9185.81/G-29/16, 35-9185.81/G-70/21, T-39-20, 35-9185.82/A-8/20) and procedures were performed to their ethical guidelines.

### Animals

Mice (male and female, 7-16 weeks of age) were housed with food and water ad libitum on a 12 h light/dark cycle (housing conditions 20-22 °C, 45-65 % humidity).

#### Mouse line

“Ntsr1-Cre-ChR2-EYFP”; crossbreed between “Ntsr1-cre” (B6.FVB(Cg)-Tg(Ntsr1-cre)GN220Gsat/Mmucd) x “Ai32” (B6;129S-Gt(ROSA)26Sortm32(CAG-COP4^*^H134R/EYFP)Hze/J)

### Histology and immunohistochemistry

The mice were humanely euthanised (i.p. Ketamin 120 mg/kg KG; Xylazin 20 mg/Kg KG) followed by transcardial perfusion using a solution of 4 % paraformaldehyde (PFA) in phosphate-buffered saline (PBS). Brain tissue was sliced into 80 µm sections using a vibratome (Thermo Scientific Microm HM 650 V) and visualised using an epifluorescent microscope (Leica DM6000).

### In vivo electrophysiology

#### Anaesthetised in vivo electrophysiology

Mice were anaesthetised using Urethane (1.4 g/kg, i.p.) and anaesthesia was maintained using an oxygen-isoflurane mixture (0.2 %). Mice were securely positioned with ear bars, and the skull was levelled. A craniotomy was performed above the well site, and a well, filled with isotonic Ringer’s solution, was cemented in place using Paladur. Silicon probes with 64 sharpened channels (impedance ∼50 kOhm) (Cambridge Neurotech) were inserted into the S1-HL cortex (ML = +1.67 mm, AP = −1.0 mm, DV = −1.4 mm) and/or VPL (ML = +1.8, AP = − 1.3, DV = −4.5) using a micromanipulator (Luigs Neumann 3-axis Motor) at a rate of approximately 2 μm/s. The probes were connected to a RHD2164 headstage amplifier chip (Intan technologies) via a connector (ASSY-77) and an adaptor (A64-Om32×2 Samtec). Signals were amplified and digitised at a sampling rate of 30,030 Hz using an RDH2000 Intan evaluation board through a USB 2.0 interface. Data acquisition was performed using an Intan Talker module (Cambridge Electronic Devices, Cambridge, UK) with Spike2 (v9.06) software.

#### Spike Sorting

Voltage data underwent band-pass filtering upon initial acquisition (0.1-10,000 Hz or 500-5000 Hz). Subsequently, Spike2 data files (.smrx) were converted into binary files through a conversion process that comprised reading the electrophysiology channels within the.smrx file, converting them back to uint16 values from the 16-bit depth analog-to-digital (ADC), and then writing this information into the resulting.bin file. Spike sorting procedures were carried out in a semi-automated manner using Matlab-based Kilosort 2.5, followed by the curation of resulting clusters in Phy2 (available at https://github.com/cortex-lab/phy). single units meeting specific criteria—having less than 0.5 % refractory period violations (within 1 ms) and a baseline spike rate greater than 0.1 Hz—were selected for further analysis. The spike-sorting parameters for Kilosort 2.5 were as follows:

| Parameter | Value | Parameter | Value |
| --- | --- | --- | --- |
| ops.fs | 3.003003003003003e+04 | ops.sigmaMask | 30 |
| ops.fshigh | 600 | ops.ThPre | 8 |
| ops.Th | [10 2] | ops.sig | 20 |
| ops.lam | 10 | ops.nblocks | 0 |
| ops.AUCsplit | 0.7 | ops.spkTh | -6 |
| ops.minFR | 1/50 | ops.reorder | 1 |
| ops.momentum | [20 400] | ops.nskip | 35 |
| ops.GPU | 1 | ops.Nfilt | 1024 |
| ops.nfilt_factor | 4 | ops.ntbuff | 64 |
| ops.NT | 64*1024+ ops.ntbuff | ops.whiteningRange | 32 |
| ops.nSkipCov | 25 | ops.scaleproc | 200 |
| ops.nPCs | 6 | ops.useRAM | 0 |

#### Putative cell-type classification

##### Thalamic units

VPL units were identified based on probe location within VPL through histological examination. The recording process involved the identification of the dorso-lateral channels containing units that exhibited significant responses to the continuous laser condition. All units within this determined range were classified as VPL units.

##### Cortical units

Recording channels from silicon probes were aligned with histological layer demarcations to attribute each unit a specific cortical depth and layer. Histological estimations of layer borders within the S1-HL cortex were determined using characteristics such as fluorescent soma sizes and densities, (refer to example in **Fig. 1b**). Units presumed to be L6CT were singled out based on their response characteristics to laser light pulses, exhibiting low latency and low jitter (see **Fig. 1d**). Waveform parameters were not used to separate units by waveforms, in this analysis (Barthó et al. 2004; Cardin et al. 2009; Madisen et al. 2012; Halassa et al. 2014; Anastassiou et al. 2015; Schmitt et al. 2017; Yu et al. 2019). Electrophysiological analyses therefore encompassed putative FS and RS units.

**Figure 1:**
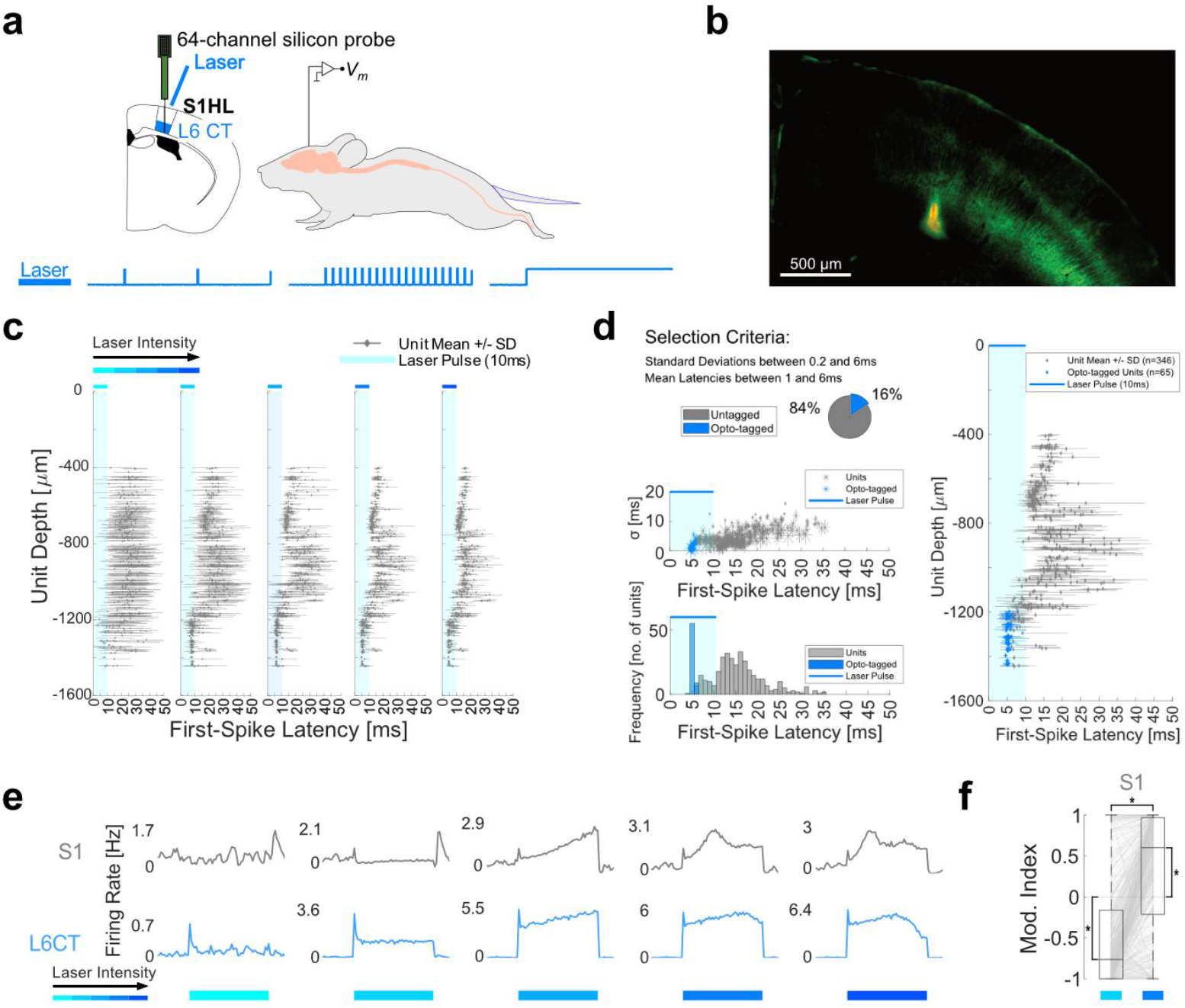
S1 L6CT bidirectionally modulates cortical activity in an activity-dependent manner. **(a)** 10 Hz laser trains (10 ms pulse length) were applied to S1-HL of Ntsr1-Cre-ChR2-EYFP mice (*n* = 5 mice) as part of a 5 seconds on 5 seconds off protocol (> 1000 pulses in total per mouse). Mean first-spike latencies and standard deviations of the first spike latencies to all 10 ms laser pulses were calculated per unit from a pooled dataset. **(b)** Example coronal section of S1 in Ntsr1-Cre-ChR2-EYFP mouse with DiI-stain (yellow) denoting the tip of the silicone probe. **(c)** single unit first-spike latencies plotted against cortical depth for increasing laser stimulation intensities (left to right: 58, 123, 382, 802, 1201 mW/mm^2^ from fibre tip). **(d)** Example L6CT optotagging (802 mW/mm^2^ from fibre tip) based on first-spike latency and standard deviation (σ) criteria (left), revealing optotagged units (blue) corresponding to the deepest units recorded (right). **(e)** Example population PSTHs of S1 (non-tagged, top) and L6CT (bottom) populations for increasing laser stimulation intensities (left to right: 58, 123, 382, 802, 1201 mW/mm^2^ from fibre tip). **(f)** Modulation indices from *n* = 5 mice showing suppression of S1 activity (791 S1 units, median MI = − 0.76) under low L6CT population activity (215 L6CT units, median L6CT low = 0.83 Hz) and increased S1 activity (791 S1 units, median MI = 0.6) under high L6CT population activity (215 L6CT units, median L6CT high = 3.77 Hz). Lines denote individual unit MI movement between conditions. ^*^ represents *p* < 0.05; 8d: Wilcoxon signed-rank test. Exact *p* values in Supplementary Table 1.

Optotagged units were verified by plotting these units based on their depth along the cortical axis, illustrating the expected positioning of optotagged L5 and L6CT neurons within layers 5 and 6, respectively (refer to **Fig. 1d**).

#### Spike train analysis

Following spike-sorting, spike times were aligned to stimulus onsets and segregated into stimulation conditions utilising customised scripts in Matlab 2022a.

##### Spike train statistical analysis

All statistical analysis was done in Matlab 2023a, using built-in or custom-written functions. For the modulation index analysis in **Figs. 1f and 2c**, the considered analysis windows were the final 2 seconds for each 5 second pulse. Baseline activity was considered as the final 2 seconds before the onset of each 5 second stimulus. Modulation index (MI) was calculated as 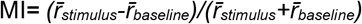 as in (Onodera and Kato 2022). High L6CT conditions were classified as laser intensity > 382 mW/mm^2^, Low L6CT as laser intensity <= 382 mW/mm^2^. Wilcoxon signed-rank tests were used for single unit comparisons between conditions, and Wilcoxon rank sum tests were used for comparisons between groups.

##### Population Synchrony Quantification

After identifying the units belonging to each subpopulation, triggered Peri-Stimulus-Time-Histograms (PSTHs) were calculated for each condition across the 5 second condition duration. This resulted in a unit-by-bin-by-trial matrix for each condition, with each time bin containing a value of spike counts. The value in each bin denotes that for a given unit during a given trial, the unit spiked n times within a given time period, with a value of 0 denoting that the unit did not spike. By transforming the matrix such that n > 0 = 1, the bins in this ‘Coactivity PSTH’ now denote whether the unit participated at all for a given time bin in a given trial. The sum of the bins across units constitutes a measure of coactivity for the population at a given time period for a given trial. The coactivity measure for a given PSTH was calculated as the mean coactivity across the bins where at least one unit during that time period was active. The coactivity measure per condition is very sensitive to how many bins each trial is partitioned into, and differs between optogenetic stimulation conditions. We leveraged this difference by plotting Coactivity PSTHs for a range of bin sizes from 0.1 ms to 2.048 seconds. The coactivity measures for each condition changed as the bin sizes increased, but the rate of change differed between conditions. This too is sensitive to the choice of bin size, so the final step was to plot the change in coactivity divided by the change in bin size to give a curve where the maximum value of the rate of change is a representation of the increase in bin size necessary to capture the greatest increase in average coactivity. The higher the score the closer a population’s neurons tend to spike together.

##### Thalamic Burst Identification

Thalamic bursts were classified as 2 or more spikes that each do not exceed 6 ms inter-spike-intervals, which are preceded by at least 50 ms prior silence, and do not exceed 100 ms total duration. These parameters were based on the ones used to study nociceptive signalling in VPL by *Hains et al*. but with a stricter inter-spike interval (ISI) threshold and a pre-burst silence to reflect prior hyperpolarisation necessary for T-Type Ca^2+^ channel availability (Zhan et al. 1999; Hains et al. 2006).

## Results

To assess how L6CT output modulates cortical population activity, we recorded multilayer activity across the S1 hind limb (HL) cortex in anaesthetised Ntsr1-Cre-Chr2-EYFP mice (*n* = 5) whilst optogenetically stimulating L6CT neurons at increasing light intensities. Stimulating L6CT at 10 Hz with increasing light intensity led to a progressive decrease in the post-pulse first-spike latencies across cortical populations, with the deepest units exhibiting the shortest latencies already at lower light intensities (**Fig. 1c**). At higher intensities, more superficial units also showed decreased latencies, though to a lesser extent than deep units.

Putative L6CT were identified through ‘optotagging’ based on their low first-spike latencies after light onset and low standard deviation in first-spike latencies. As expected, these units were localized to the deepest cortical electrodes, corresponding to layer 6 (**Fig. 1d**). Separating the L6CT population from the rest of the S1 population, allowed us to examine how different levels of L6CT activity influence S1 firing. At the lowest laser intensity, insufficient to sustain elevated L6CT activity, S1 activity remained unchanged. However, at sustained yet low L6CT activity, S1 activity was suppressed throughout the period of optogenetic stimulation (**Fig. 1e**). Surprisingly, increasing L6CT activity further caused a reversal of this effect, with S1 firing rates progressively rising above baseline and peaking earlier as a function of L6CT activity.

To assess whether this population-level shift reflected heterogeneous responses across S1 subpopulations, we calculated modulation indices (MI) to track individual unit responses across different L6CT activity conditions. While low L6CT activity globally reduced MI across the S1 population, high L6CT activity globally increased MI (**Fig. 1f**). Notably, 65% (517/791) of S1 units exhibited an increased MI from low to high L6CT activity, with ∼40% (316/791) switching from suppression to excitation. Only ∼10% (84/791) showed reduced MI at high L6CT levels compared to low L6CT. These results demonstrate that sustained L6CT activity bidirectionally modulates cortical excitability as a function of L6CT firing rate, transitioning from S1 suppression at low L6CT firing rate to S1 excitation at higher L6CT firing rates.

The laser response profiles shown in **Fig. 1 c & d** suggest three subpopulations that are separable by their respective first-spike latencies and depths. Analysis of first-spike latencies and unit depths revealed three distinct subpopulations within S1, corresponding to putative layer 4 (L4), putative layer 5 (L5) and L6CT neurons. L4 units had longer mean first-spike latencies with lower variability compared to L6CT units and resided on the upper recording channels, whereas L5 units resided between L6CT and L4 and had the longest mean first-spike latency with the highest variability (**Fig. 2a**). L5 units also had a higher basal firing rate than L4 and L6 populations, (median L4 = 0.09 Hz, interquartile range (IQR) = 0.27; L5 = 0.51 Hz, IQR = 1.24; L6CT = 0.02 Hz, IQR = 0.23).

**Figure 2:**
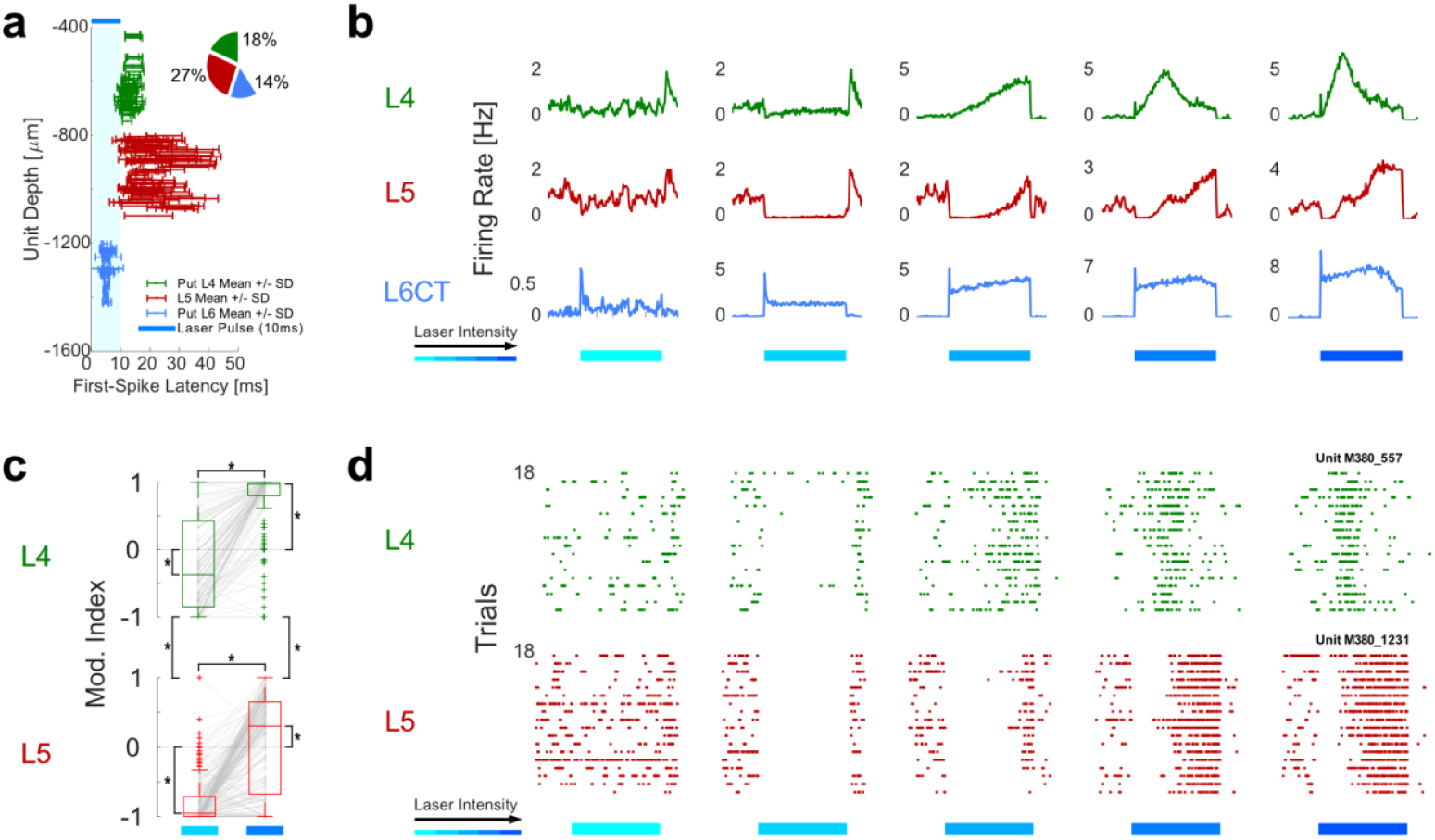
Cortical subpopulations exhibit distinct L6CT-mediated activity switches. **(a)** Example latency vs depth plot (from one recording) with three populations grouped according to their first spike latency and variability in response to L6CT laser stimulation. L6CT units (58 units, blue), putative layer 4 (75 units, green) and putative layer 5 (112 units, red) located at different cortical depths. **(b)** Example population activity in response to increasing L6CT stimulation intensity for putative L4 (green), L5 (red) and L6CT (blue). **(c)** Box plots showing modulation indices for putative L4 (green) and L5 (red) under lower (light blue) and higher (dark blue) L6CT activity (median L6CT low = 0.83 Hz, median L6CT high = 3.77 Hz). Low L6CT activity induced significantly lower L4 activity (163 units, median MI = −0.38) as well as L5 suppression (219 units, median MI = −0.95). In contrast, high L6CT activity increased L4 (median MI = 0.98) and L5 activity (median MI = 0.3). **(d)** single unit raster plots showing putative L4 (green) and L5 (red) single unit responses to increasing light intensities of L6CT stimulation. ^*^represents *p* < 0.05; 2c: Wilcoxon signed-rank test (within-group differences), Wilcoxon rank sum (between-group differences). Exact *p* values in Supplementary Table 1.

Both, L5 and L4 units exhibited suppression during low L6CT activity, though they differed in the rate at which increasing L6CT activity switched their activity from suppressed to excited (**Fig. 2b**). Unit-by-unit quantification of this suppression revealed a down-modulation of both L4 (*n* = 5 mice, 163 units) and L5 (*n* = 5 mice, 218 units) (**Fig. 2c**). However, the magnitude of suppression differed between populations, with L5 units showing a greater reduction (MI; median MI = −0.96) compared to L4 units (median MI = −0.31).

In the high L6CT condition, both L4 and L5 exhibited increased activity that differed significantly between each other, with L4 (median MI = 0.98) showing greater enhancement than L5 (median MI = 0.30). Just over half of the L4 units (82/163) switched their MI sign from low to high L6CT activity, with 76 % (124/163) showing an increased MI. Similarly, about half of the L5 units (119/218) switched their MI sign from low to high L6CT activity, with 78 % (169/218) showing an increased MI. These findings indicate that L6CT-mediated bidirectional modulation affects cortical subpopulations differently, with the extent of suppression and excitation depending on cortical layer (**Fig. 2d**).

The bidirectional activity switch of L4 and L5 populations in response to increased L6CT activity indicates non-linear responses to increased L6CT spiking. As L6CT activity is increased, the probability of convergent inputs arriving to trigger action potentials may also increase and could therefore be reflected in a greater spiking probability across the L4 and L5 subpopulations. The L6-L4 synapse has previously been demonstrated *in vitro* to exhibit paired-pulse facilitation (Lee and Sherman 2008).

To test whether L6CT induces frequency-dependent facilitation in L4 and L5 populations we stimulated L6CT with 10 ms light pulses at different frequencies but constant laser strength (**Fig. 3a**). L4 units exhibited frequency-dependent facilitation, with higher L6CT stimulation frequencies eliciting greater activity relative to baseline (**Fig. 3b**). In contrast, L5 units showed suppression at low L6CT frequencies for at least 50 ms after the light stimulus onset. As the frequency of L6CT stimulation increased, the length of L5 suppression decreased and turned into increased L5 activity, yielding a net increase in L5 firing rates in the 50 ms following L6CT stimulation, although an initial inhibition remained. To directly assess whether this facilitation is frequency-dependent, we compared responses in the 50 ms following L6CT stimulation between the 1st and 3rd pulses in both the 1 Hz and 10 Hz conditions (**Fig. 3c**). L4 units increased firing to the 3rd stimulus in both conditions, with greater facilitation at 10 Hz. In contrast, L5 units increased firing only in response to the 3rd pulse in the 10 Hz condition (**Fig. 3d**). When comparing the change in responses between 1st and 3rd pulses across conditions, 10 Hz induced a larger increase in MI vs 1 Hz in both L4 and L5. Notably, when compared to continuous L6CT stimulation, which induced net suppression in most L4 and L5 units, 10 Hz stimulation facilitated responses. (**Fig. 3e**). Thus, L4 and L5 can be facilitated or suppressed depending on the nature of L6CT activity.

**Figure 3:**
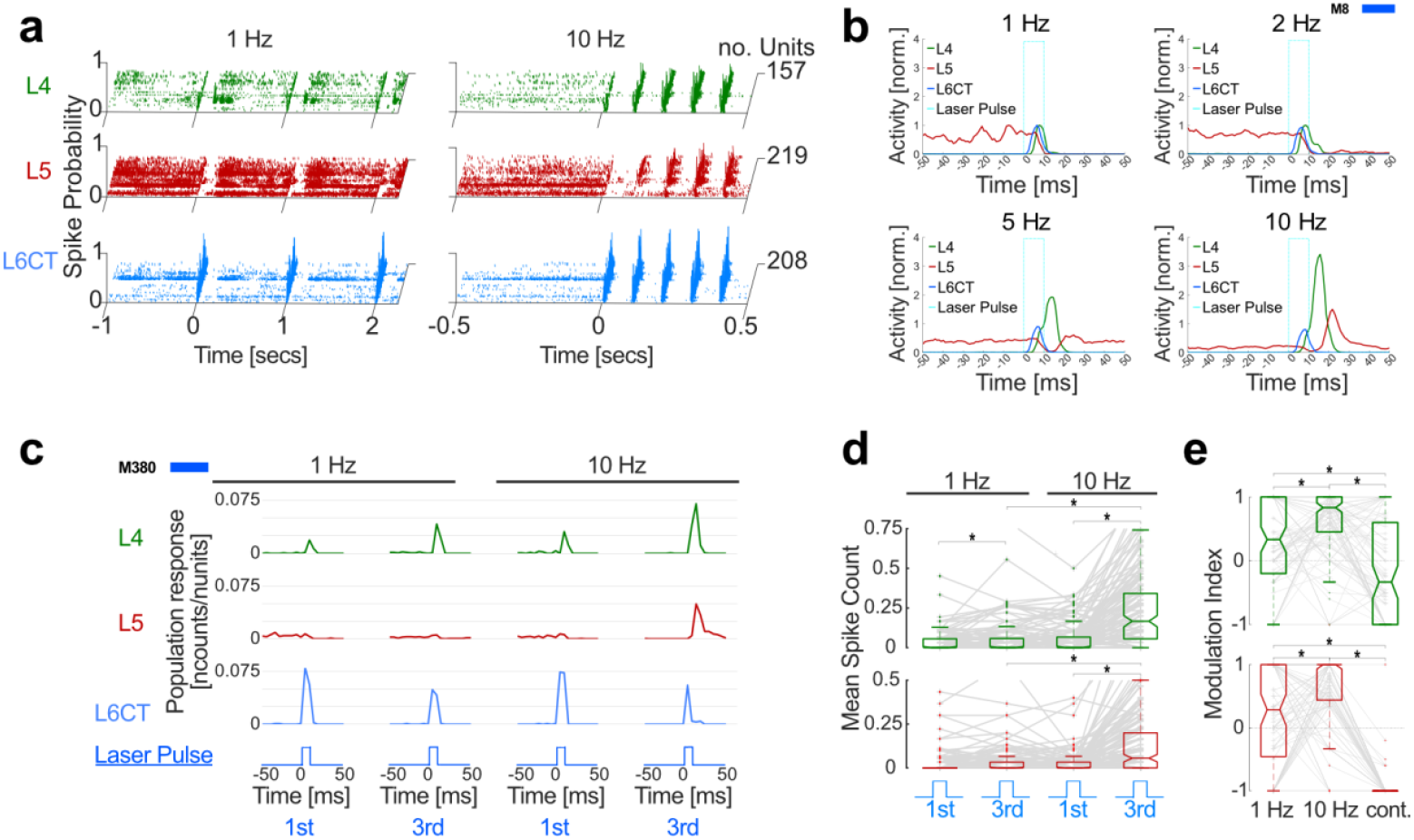
L6CT excitation causes frequency-dependent facilitation in L4 and L5. **(a)** Unit PSTHs across trials for 1 Hz (left) and 10 Hz (right) stimulation of L6CT units (blue) and corresponding responses of L5 (red) and L4 (green) units, *n* = 5 mice. Spike probability of 1 corresponds to 1 spike per trial per bin (1ms). **(b)** Example population PSTHs of L6CT high stimulation (laser pulse: light blue dotted line) for 1 Hz (top left), 2 Hz (top right), 5 Hz (bottom left) and 10 Hz (bottom right) frequencies. Normalised with respect to max. subpopulation response to 1Hz stimulus. Units from a representative experiment: L4 = 15, L5 = 31, L6CT = 39. **(c)** Example PSTHs (5 ms bins) of L6CT, L4 and L5 population responses to the 1st and 3rd pulse (10 ms) of the L6CT stimulation in the 1 Hz and 10 Hz condition. Data from one representative experiment (unit counts: L4 = 75, L5 = 112, L6CT = 58). **(d)** Mean spike counts for L4 (green) and L5 (red) units in response to the 1st and 3rd of the L6CT stimulation in the 1 Hz and 10 Hz condition. Pooled from 5 experiments: L4 = 157, L5 = 219, L6CT = 208. **(e)** Modulation indices for responses to the 1st and 3rd pulse of L6CT stimulation in the 1 Hz and 10 Hz conditions and in response to continuous pulses (cont.). Response windows were matched across all three conditions. Pooled from n = 5 mice (unit counts: L4 = 157, L5 = 219, L6CT = 208. ^*^represents *p* < 0.05; 3d & e: Wilcoxon signed-rank test. Exact *p* values in Supplementary Table 1.

Given the contrasting effects of cortical responses to 10 Hz vs continuous L6CT stimulation, (**Fig. 4a & b**), we asked whether L6CT synchrony, rather than overall L6CT activity, might be a key determinant of L4 and L5 responses. To quantify synchrony independently of firing rates, we binned the spiking activity during the response windows into PSTHs of varying bin sizes. We then calculated the average population coactivity as the number of spiking units in a given bin for a given trial over a range of bin sizes. Because coactivity depends on bin size (larger bins tend to contain activity from more units), we measured how rapidly coactivity changes as a function of bin size. The largest differences between conditions emerged over bin size increases within 5 ms and the maximum value was taken as the measure of population synchrony (**Fig. 4c**).

**Figure 4:**
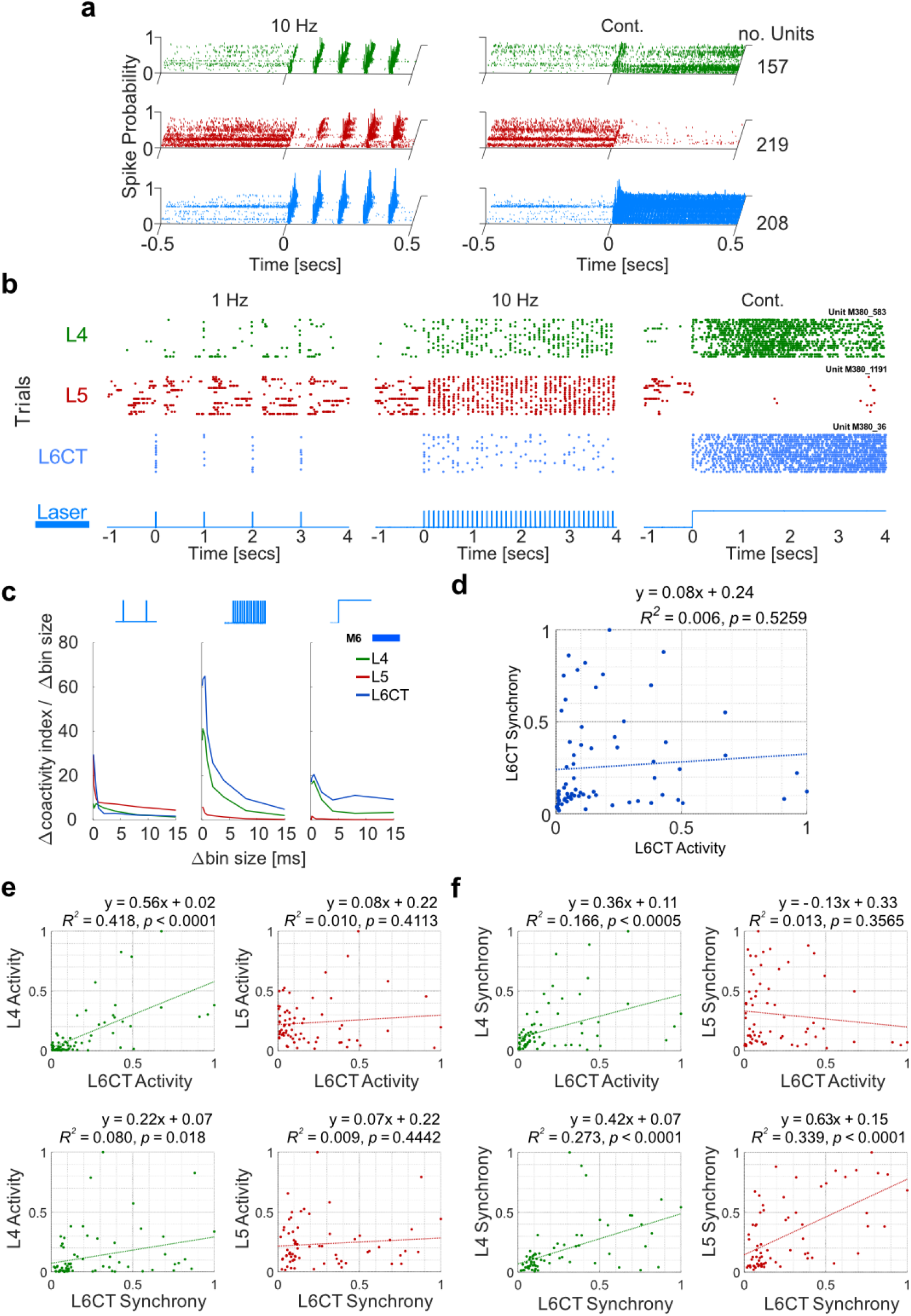
L6CT entrains L4 and L5 by influencing population firing synchrony. **(a)** Unit PSTHs for 10 Hz (left) and continuous (right) stimulation of L6CT (blue) and corresponding responses in L4 (green) and L5 (red) across trials. *n* = 5 mice. Spike probability of 1 = 1 spike per trial per bin for a given unit. Bin size = 1 ms. **(b)** Example unit raster plots displaying responses of L6CT, L4 and L5 to optogenetic stimulation under different conditions (1 Hz, 10 Hz, continuous). **(c)** Example synchrony plot. N units: L4 = 20, L5 = 15, L6 = 29. Synchrony was calculated as the maximum increase in the number of coactive units as a function of increasing bin size (The peak y-value on each graph), normalised to the max of all stimulation conditions across all experiments (*n* = 70). **(d)** Normalised Activity vs Synchrony scatter plot for L6CT pooled across experiments demonstrating little covariance between measures, indicating sufficient extraction of independent measures of synchrony and activity. **(e)** Effect of L6CT activity (top) and synchrony (bottom) on activity in L4 (green) and L5 (red). L6CT activity predicts > 40% of L4 activity variability. **(f)** Effect of L6CT activity (top) and synchrony (bottom) on synchrony in L4 (green) and L5 (red). A-d calculated with response windows 0-5 seconds. B-d: 70 data points across 5 experiments.

L6CT synchrony and overall activity were effectively uncorrelated across conditions (slope = 0.08; **Fig. 4d**), indicating that these are distinct and independently varying parameters. At high laser intensities, the low frequency stimulation conditions produced highly synchronous but low L6CT activity, whereas continuous stimulation had the opposite effect (**supp. Fig. 1a**). When examining the influence of L6CT activity and synchrony on the other cortical populations, L6CT activity predicted L4 activity but not L5 activity (**Fig. 4e**). Neither L4 activity nor L5 activity were correlated significantly with L6CT synchrony. In contrast, L6CT synchrony significantly predicted L4 and L5 synchrony, and L4 synchrony was further correlated significantly with L6CT activity (**Fig. 4f**).

Taken together, L6CT spiking exerts a strong influence on L4 spiking, with its activity shaping both L4 activity and synchrony. Additionally, L6CT entrains both L4 and L5 spiking through its influence on population spiking synchrony whilst in the case of L5 not necessarily altering overall activity. Intriguingly, L5 activity was significantly correlated with L4 activity (**supp. Fig. 1b**). Combined with the observed temporal sequence of activation (L6CT → L4 → L5; Fig. **2a**, **3b**), this suggests that L6CT drives L4 activity and synchrony, which in turn influences L5 firing.

After establishing L6CT’s impact on cortical dynamics, we next tested how different L6CT spiking patterns modulate thalamic. We recorded from VPL in Ntsr1-Cre-Ch2-EYFP mice during optogenetic stimulation of L6CT neurons. In one instance, dual recording in S1 and VPL was achieved to observe activity in both L6CT and VPL populations simultaneously (**Fig. 5a-c**).

**Figure 5:**
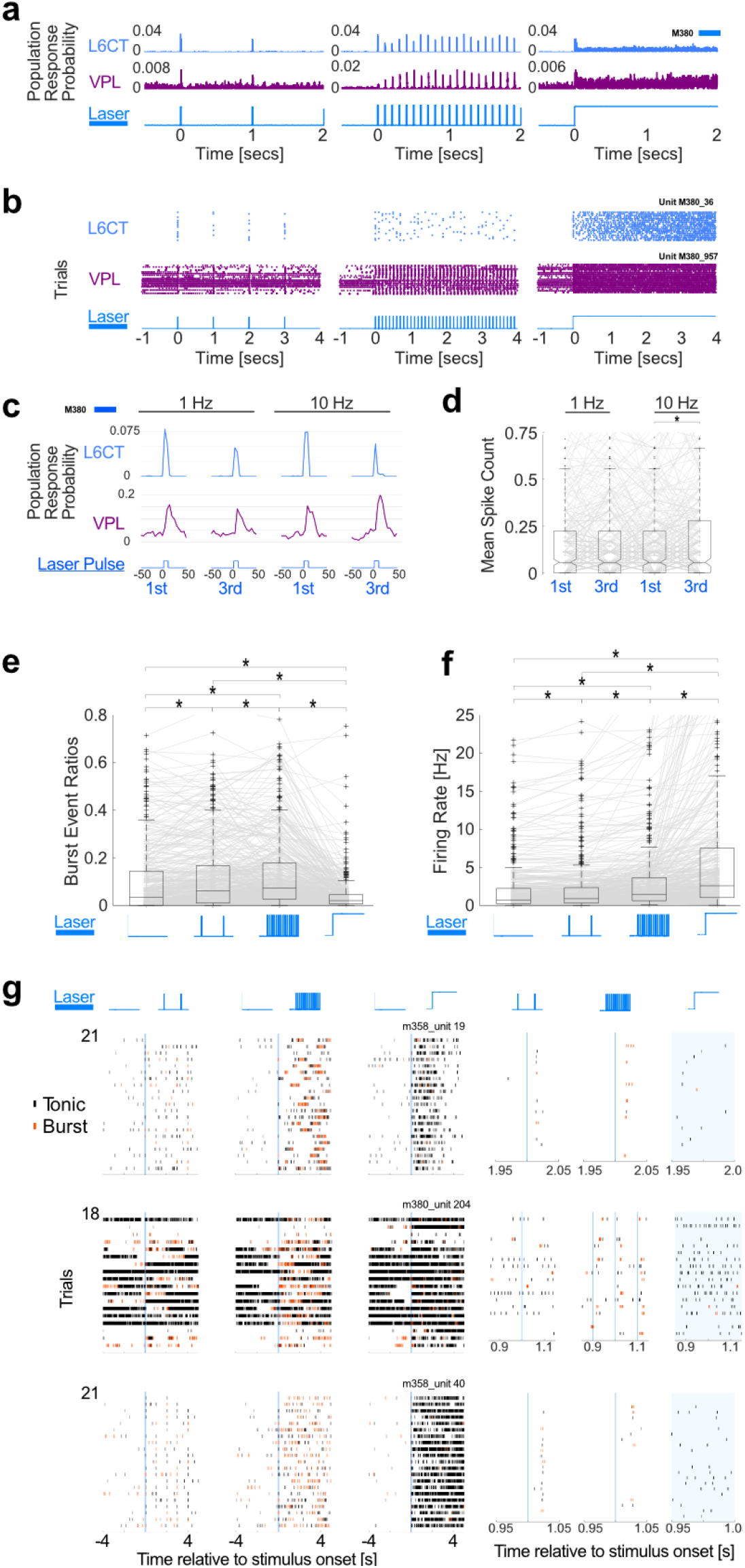
L6CT stimulation patterns shape VPL spike rates and firing modes. **(a)** Example Population PSTHs for L6CT (top) and VPL (middle) populations simultaneously recorded during 1 Hz (left), 10 Hz (middle), and continuous (right) optogenetic stimulation of L6CT neurons. Viewing window shortened (−0.5 - 2 seconds around onset) to illustrate short-term dynamics. **(b)** Example Raster plots of L6CT (top) and VPL (middle) single units simultaneously recorded during 1 Hz (left), 10 Hz (middle), and continuous (right) optogenetic stimulation of L6CT neurons. **(c)** Example dual recording in S1 and VPL demonstrating increased VPL activity to the 3rd pulse of 10 Hz condition. **(d)** VPL population shows increased response to 3rd pulse of a 10 Hz train, but not 3rd pulse of a 1 Hz train. 439 units, *n* = 4 mice. **(e)** Frequency stimulation increased burst event ratios relative to baseline in a frequency-dependent manner (10 Hz > 1 Hz), whereas continuous stimulation decreased burst event ratios relative to baseline. 439 units, *n* = 4 mice. **(f)** L6CT stimulation increased total firing rate relative to baseline, displaying a dose-response relationship with respect to **(g)** Example Raster plots of VPL single units to 1 Hz, 10 Hz, and continuous L6T optogenetic stimulation. Spikes coloured in orange belong to burst events, those in black do not belong to bursts. Panels to the right are enlarged sections of those on the left. ^*^represents *p* < 0.05; 5d & f-h: Wilcoxon signed-rank test.Exact *p* values in Supplementary Table 1.

The L6-CT synapses in the ventroposterior thalamic nucleus and lateral geniculate nucleus have previously been shown to undergo paired-pulse facilitation (Jurgens et al. 2012; Crandall et al. 2015; Kirchgessner et al. 2020). To address this in the VPL, we measured VPL spiking responses following the 1st and 3rd optogenetic L6CT stimulation pulses in 1 Hz and 10 Hz conditions (**Fig. 5c**). Consistent with facilitation, 10 Hz – but not 1 Hz stimulation, facilitated VPL spike output (**Fig. 5d**).

Previous studies have demonstrated that continuous L6CT excitation induces a switch in neurons from burst to tonic mode (Mease et al. 2014; Ziegler et al. 2023). However, the effects of different L6CT stimulation frequencies on thalamic firing modes remain less explored. We compared 1 Hz, 10 Hz, and continuous L6CT stimulation on VPL firing. Burst spikes were defined as ≥ 2 spikes with ≤ 6 ms inter-spike intervals, preceded by ≥50 ms silence, and lasting ≤100 ms (**Fig. 5e**). As reported in previous literature, we found that continuous stimulation of L6CT indeed decreased VPL bursting, measured as the proportion of bursts spikes to all spikes, indicating a shift towards tonic-firing mode (**Fig. 5f**). In contrast, both 1 Hz and 10 Hz L6CT stimulation increased bursting relative to baseline, with 10 Hz producing the strongest effect.

All L6CT stimulation patterns elevated overall VPL firing rates, in a manner proportional to L6CT activity, with 1 Hz eliciting the smallest increase in spike rate and continuous stimulation the largest (**Fig. 5h**). These findings suggest that L6CT excitation enhances thalamic throughput in a manner proportional to the L6CT activity level, while the temporal pattern of L6CT spiking determines whether VPL neurons engage burst or tonic firing modes. total optogenetic stimulation. 439 units, *n* = 4 mice.

## Discussion

Whether corticothalamic layer-6 (L6CT) activity suppresses or enhances cortical firing has remained contentious, with studies across sensory cortices reporting either an inhibitory or excitatory effect of optogenetic L6CT excitation, but not both within the same sensory cortex (Olsen et al. 2012; Guo et al. 2017; Ziegler et al. 2023; Dimwamwa et al. 2024). In a similar approach to prior studies (Kvitsiani et al. 2013; Hangya et al. 2015; Ziegler et al. 2023), we used an optotagging procedure based on the first-spike latencies to repeated pulses of optogenetic stimulation and their standard deviations, which enabled separation of S1 L6CT single units from the rest of the recorded S1 single units. Here we demonstrated a switch in the activity of the same population in S1 cortex that depended on the level of optogenetic L6CT excitation, with lower L6CT activity suppressing basal activity in S1 layers 4 and 5, whereas higher L6CT excitation facilitated activity in these layers. Rather than this activity switch resulting from heterogeneously behaving populations within each layer, where one subpopulation is suppressed under low L6CT activity and the other facilitated under high L6CT activity, the activity switch was carried by single units, with approximately half of the units switching from suppressed to elevated activity as L6CT drive increases. Whilst previous studies in awake and anaesthetised mice have suggested a mainly inhibitory role of L6CT in S1 (Ziegler et al. 2023; Dimwamwa et al. 2024), we demonstrate that S1 L6CT can exert either an inhibitory or excitatory effect on cortical activity dependent on the dynamics of L6CT activity. This raises the question of under what behavioural circumstances L6CT is active, and to what extent. It furthermore potentially implicates L6CT as a context-dependent suppressor or booster of sensory computation in S1 and possibly other sensory cortical regions. L6CT modulated two distinct cortical subpopulations in both directions, with L4 preferentially excited, and L5 preferentially inhibited by L6CT activity. These subpopulation-dependent variations in bidirectional modulation by L6CT indicates that this switch can occur at different types of synapses and is likely not narrowly limited to finely tuned synaptic and circuit configurations. Therefore, L6CT-cortical circuitry is unlikely to be a unique example of a bidirectional switch based on activity level and could be a feature of many pathways. Indeed, a similar bidirectional switch has been observed in thalamic circuits to varying degrees of L6CT activity (Crandall et al. 2015). Urethane is reported to suppress spiking activity in cortical pyramidal neurons (Sceniak and Maciver 2006; Potez and Larkum 2008), suggesting that at higher baseline activity of L6CT under awake conditions, the system is set towards L6CT facilitation.

L6CT-evoked spikes in downstream neurons had longer latencies and variability in L5 than L4, suggesting that L5 spikes arise through multiple synapses. This is corroborated by population PSTH analysis showing initial L5 inhibition followed by rebound elevated activity above baseline. In agreement with previous results from hind limb S1 (Ziegler et al. 2023), L6CT powerfully suppresses L5 and may subsequently receive sufficient excitation through an intermediary. That L4 activity was a much stronger predictor of L5 activity than L6CT activity may also indicate that L4 is an intermediate cause of L5 excitatory responses under high L6CT activity. We observed that L4 also undergoes frequency-dependent facilitation to L6CT activity, which could partially explain the frequency-dependent L5 facilitation observed, through an increased excitatory drive. We observed a notably robust positive correlation between L6CT activity and L4 activity, as well as frequency-dependent facilitation in L4 units to repeated pulses of L6CT excitation, possibly due to implied direct facilitatory effects overcoming metabotropic suppressive modulation (Lee and Sherman 2008, 2009, 2012; Kim et al. 2014). This frequency-dependent facilitation of L4 could aid in amplifying sensory inputs from thalamic regions, of which a substantial proportion terminates in L4 (Gilbert and Wiesel 1979; Sherman and Guillery 2002; Douglas and Martin 2004; Clascá et al. 2012; Constantinople and Bruno 2013; Adesnik and Naka 2018). Another important finding was the facilitation observed in L4 to 1 Hz L6CT stimulation, which was absent in L5, indicating that L4 may integrate information from L6CT over much longer time windows than L5 does; in essence L4 has a longer ‘memory’ of L6CT activity compared with L5.

Although optogenetic stimulation is typified by simultaneous excitation of opsin-expressing neurons (Boyden et al. 2005; Han et al. 2009), we developed a methodological tool to quantify the relative change in synchrony of neuronal spiking between different optogenetic stimulation conditions to investigate whether changes in synchrony within a range of already somewhat synchronous L6CT activity resulted in changes in the synchrony and/or activity of neuronal spiking in S1. This synchrony measure did not correlate with relative changes in activity over the range of stimulation protocols delivered, and therefore allowed us to ask whether changing the relative synchrony of the L6CT spiking resulted in changes in the dynamics of downstream populations of single units in L4 and L5, independent of changes in L6CT activity. Similarly, whilst urethane anaesthesia is reported to increase slow oscillatory activity, with Up states of synchronous spiking activity (Clement et al. 2008), by cycling through conditions instead of applying conditions sequentially we were able to assess how different patterns of L6CT activity altered S1 activity and synchrony despite possible slow changes to oscillatory behaviour from the anaesthesia.

Whilst L6CT activity was a strong predictor of L4 activity, that in turn influenced and preceded L5 activity, L6CT activity did not strongly predict L5 activity. However, L6CT synchrony was a strong predictor of both L4 and L5 synchrony. Thus, L6CT appears to control the timing of L5 spiking across L5 units, but not necessarily the magnitude. If there is information transfer along L6CT→L4→L5, it would appear that L6CT neuronal synchrony is preserved more robustly than its total spiking output, and L6CT synchrony and activity could represent distinct channels during information transfer (Buzsáki 2010; Buzsáki and Vöröslakos 2023). Our research suggests that L6CT is able to impart its synchrony to other cortical layers, distinct from its effects on global cortical firing rates. It has been previously demonstrated that L6CT continuous stimulation can induce translaminar modulation in the cortex, but these findings suggest L6CT at physiological frequencies can entrain cortical spiking in upper layers to specific L6CT rhythms (Olsen et al. 2012; Guo et al. 2017; Adesnik 2018).

On the thalamic level, L6CT exerts a complementary, frequency-dependent control over VPL firing mode. Rhythmic L6CT input (1 or 10 Hz) increased bursting relative to baseline, whereas continuous L6CT activity shifted thalamic activity towards tonic firing. This was in contrast to VPL overall firing rate, which increased as a function of total L6CT stimulation duration. This suggests that VPL throughput may increase with progressively greater L6CT activity, but VPL burst probability may be altered within a somewhat independent range, in a non-linear fashion dependent on the rhythmicity or synchrony of L6CT activity, potentially representing distinct channels of neural information transfer (Payeur et al. 2021).

Our findings converge with those of (Dimwamwa et al. 2024), who showed in awake mice that S1 L6CT activation induces bidirectional modulation of firing rates in the ventral posteromedial nucleus (VPm) of the whisker thalamus, with the direction (suppression vs enhancement) determined by L6CT firing rate and synchrony. In our anaesthetised preparations, we further demonstrate that L6CT activation not only modulates the magnitude of thalamic output but also governs the mode of thalamic firing (burst vs tonic), a key determinant of sensory signal transmission.

Taken together, this research suggests that L6CT can dynamically ‘tune up’ and ‘tune down’ signalling in S1 and can independently shape cortical synchrony, whilst altering incoming information from the thalamus via bidirectional control of thalamic firing mode.

## Acknowledgments

We thank Katharina Ziegler for helpful comments on the manuscript. The funders had no role in study design, data collection and analysis, decision to publish, or preparation of the manuscript.

## Author contributions

Conceptualization: RF, EIC, AG

Methodology: RF, EIC

Investigation: RF

Data analysis and analysis tools: RF, EIC

Visualization: RF

Funding acquisition: AG

Project administration: AG

Supervision: AG

Writing – review & editing: RF, AG

## Supplementary Materials

**Supplementary Figure S1:**
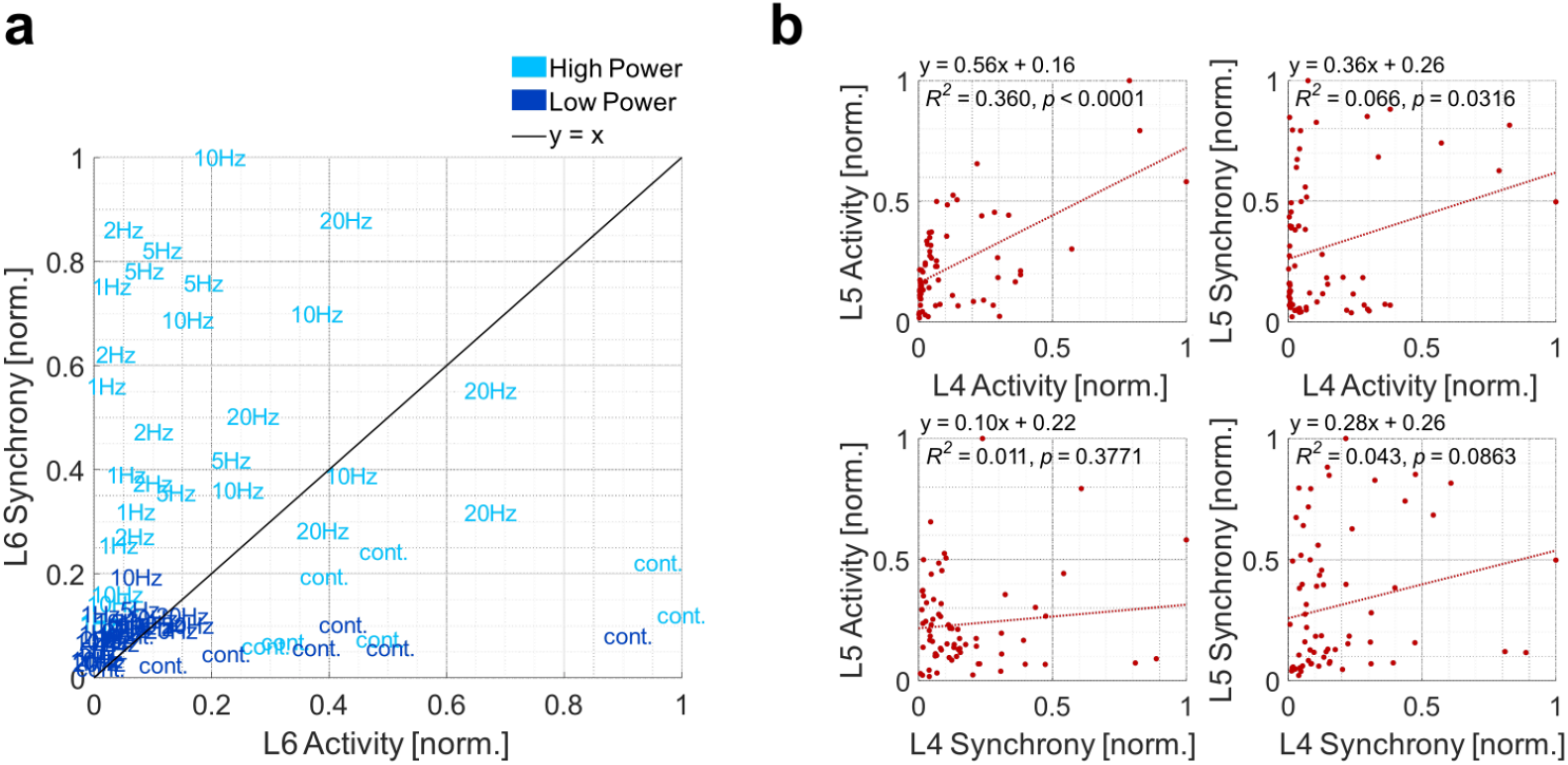
Effect of different optogenetic stimulation conditions on Synchrony-Activity Ratios, and effect of L4 on L5. **(a)** Activity vs synchrony plot for the different stimulation conditions and light intensities (High Power > 382mW/mm^2^, Low Power <= 382mW/mm^2^) for tagged L6CT populations from *n* = 5 mice (Data points are identical as for **Fig. 4d**). **(b)** Effect of L4 activity (top) and synchrony (bottom) on activity (left) and synchrony (right) in L5 (red). L4 activity predicts >35% of L5 activity variability. *n* = 5 mice.

**Supplementary Table S1.** Description of statistical parameters and *p*-values by figure.

|  |  |  |  |
| --- | --- | --- | --- |
| 1d | Low L6CT<br>Spontaneous vs<br>Evoked | $p = 7.1 \times 10^{-38}$ | Wilcoxon<br>signed-rank |
| | High L6CT<br>Spontaneous vs<br>Evoked | $p = 1.07 \times 10^{-38}$ | Wilcoxon<br>signed-rank |
| | Low L6CT vs High<br>L6CT Modulation<br>Indices | $p = 2.13 \times 10^{-76}$ | Wilcoxon<br>signed-rank |
| | L4 vs L5 Basal<br>Firing rates | Low L6CT; $p = 2.8 \times 10^{-15}$<br>High L6CT; $p = 2.6 \times 10^{-22}$ | Wilcoxon rank<br>sum |
| | L4 vs L6CT Basal<br>Firing rates | Low L6CT; $p = 1.7 \times 10^{-4}$<br>High L6CT; $p = 0.0025$ | Wilcoxon rank<br>sum |
| | L5 vs L6CT Basal<br>Firing rates | Low L6CT; $p = 4.0 \times 10^{-29}$<br>High L6CT; $p = 8.7 \times 10^{-34}$ | Wilcoxon rank<br>sum |
| 2c | Low L6CT<br>Spontaneous vs<br>Evoked | L4; $p = 8.4 \times 10^{-5}$<br>L5; $p = 9.6 \times 10^{-35}$ | Wilcoxon<br>signed-rank |
| | High L6CT<br>Spontaneous vs<br>Evoked | L4; $p = 1.2 \times 10^{-24}$<br>L5; $p = 1.8 \times 10^{-4}$ | Wilcoxon<br>signed-rank |
| | Low L6CT vs High<br>L6CT Modulation<br>Indices | L4; $p = 2.0 \times 10^{-22}$<br>L5; $p = 1.5 \times 10^{-28}$ | Wilcoxon<br>signed-rank |
| | L4 vs L5<br>Modulation indices | Low L6CT; $p = 5.2 \times 10^{-16}$<br>High L6CT; $p = 1.2 \times 10^{-29}$ | Wilcoxon rank<br>sum |
| 3d | L4 Counts<br>Facilitation | 1 Hz 1st vs 3rd pulse; $p = 5.16 \times 10^{-4}$<br>10 Hz 1st vs 3rd pulse; $p = 1.6 \times 10^{-19}$<br>1 Hz 3rd vs 10 Hz 1st pulse; $p = 0.75$<br>1st pulse 1 Hz vs 10 Hz; $p = 5.07 \times 10^{-4}$ | Wilcoxon<br>signed-rank |
| | | 3rd pulse 1 Hz vs 10 Hz; $p = 1.49 \times 10^{-18}$ | |
| | L5 Counts Facilitation | 1 Hz 1st vs 3rd pulse; $p = 0.0353$<br>10 Hz 1st vs 3rd pulse; $p = 2.98 \times 10^{-21}$<br>1 Hz 3rd vs 10 Hz 1st pulse; $p = 0.14$<br>1st pulse 1 Hz vs 10 Hz; $p = 0.59$<br>3rd pulse 1 Hz vs 10 Hz; $p = 9.09 \times 10^{-19}$ | Wilcoxon signed-rank |
| 3e | L4 Modulation Indices | 1 Hz vs 10 Hz; $p = 0.0466$<br>1 Hz vs continuous; $p = 9.5 \times 10^{-5}$<br>10 Hz vs continuous; $p = 2.7 \times 10^{-8}$ | Wilcoxon signed-rank |
| | L5 Modulation Indices | 1 Hz vs 10 Hz; $p = 0.0177$<br>1 Hz vs continuous; $p = 1.56 \times 10^{-7}$<br>10 Hz vs continuous; $p = 8.9 \times 10^{-11}$ | Wilcoxon signed-rank |
| 5d | VPL Counts Facilitation | 1 Hz 1st vs 3rd pulse; $p = 0.0757$<br>10 Hz 1st vs 3rd pulse; $p = 0.007$<br>1 Hz 3rd vs 10 Hz 1st pulse; $p = 0.50$<br>1st pulse 1 Hz vs 10 Hz; $p = 0.24$<br>3rd pulse 1 Hz vs 10 Hz; $p = 0.019$ | Wilcoxon signed-rank |
| 5f | VPL Burst Event Ratios | Baseline vs 1 Hz; $p = 5.2 \times 10^{-15}$<br>Baseline vs 10 Hz; $p = 4.7 \times 10^{-16}$<br>Baseline vs continuous; $p = 4.3 \times 10^{-17}$<br>1 Hz vs 10 Hz; $p = 0.00025$<br>1 Hz vs continuous; $p = 1.54 \times 10^{-34}$<br>10 Hz vs continuous; $p = 1.56 \times 10^{-49}$ | Wilcoxon signed-rank |
| 5g | VPL Total Bursts | Baseline vs 1 Hz; $p = 2.9 \times 10^{-39}$<br>Baseline vs 10 Hz; $p = 1.1 \times 10^{-40}$<br>Baseline vs continuous; $p = 5.4 \times 10^{-11}$<br>1 Hz vs 10 Hz; $p = 6.9 \times 10^{-15}$<br>1 Hz vs continuous; $p = 0.62$<br>10 Hz vs continuous; $p = 8.1 \times 10^{-13}$ | Wilcoxon signed-rank |
| 5h | VPL Median Firing Rates | Baseline vs 1 Hz; $p = 8.4 \times 10^{-17}$<br>Baseline vs 10 Hz; $p = 2.5 \times 10^{-38}$<br>Baseline vs continuous; $p = 1.5 \times 10^{-53}$<br>1 Hz vs 10 Hz; $p = 3.8 \times 10^{-31}$<br>1 Hz vs continuous; $p = 8.7 \times 10^{-18}$<br>10 Hz vs continuous; $p = 7.5 \times 10^{-41}$ | Wilcoxon signed-rank |

## Bibliography

Adesnik H. 2018. Layer-specific excitation/inhibition balances during neuronal synchronization in the visual cortex. J Physiol. 596:1639–1657.

Adesnik H, Naka A. 2018. Cracking the Function of Layers in the Sensory Cortex. Neuron. 100:1028–1043.

Anastassiou CA, Perin R, Buzsáki G, Markram H, Koch C. 2015. Cell type-and activity-dependent extracellular correlates of intracellular spiking. J Neurophysiol. 114:608– 623.

Barthó P, Hirase H, Monconduit L, Zugaro M, Harris KD, Buzsáki G. 2004. Characterization of neocortical principal cells and interneurons by network interactions and extracellular features. J Neurophysiol. 92:600–608.

Bortone DS, Olsen SR, Scanziani M. 2014. Translaminar inhibitory cells recruited by layer 6 corticothalamic neurons suppress visual cortex. Neuron. 82:474–485.

Boyden ES, Zhang F, Bamberg E, Nagel G, Deisseroth K. 2005. Millisecond-timescale, genetically targeted optical control of neural activity. Nat Neurosci. 8:1263–1268.

Buzsáki G. 2010. Neural syntax: cell assemblies, synapsembles, and readers. Neuron. 68:362–385.

Buzsáki G, Vöröslakos M. 2023. Brain rhythms have come of age. Neuron. 111:922–926.

Cardin JA, Carlén M, Meletis K, Knoblich U, Zhang F, Deisseroth K, Tsai L-H, Moore CI. 2009. Driving fast-spiking cells induces gamma rhythm and controls sensory responses. Nature. 459:663–667.

Clascá F, Rubio-Garrido P, Jabaudon D. 2012. Unveiling the diversity of thalamocortical neuron subtypes. Eur J Neurosci. 35:1524–1532.

Clement EA, Richard A, Thwaites M, Ailon J, Peters S, Dickson CT. 2008. Cyclic and sleep-like spontaneous alternations of brain state under urethane anaesthesia. PLoS One. 3:e2004.

Constantinople CM, Bruno RM. 2013. Deep cortical layers are activated directly by thalamus. Science. 340:1591–1594.

Crandall SR, Cruikshank SJ, Connors BW. 2015. A corticothalamic switch: controlling the thalamus with dynamic synapses. Neuron. 86:768–782.

Dimwamwa ED, Pala A, Chundru V, Wright NC, Stanley GB. 2024. Dynamic corticothalamic modulation of the somatosensory thalamocortical circuit during wakefulness. Nat Commun. 15:3529.

Douglas RJ, Martin KAC. 2004. Neuronal circuits of the neocortex. Annu Rev Neurosci. 27:419–451.

Frandolig JE, Matney CJ, Lee K, Kim J, Chevée M, Kim S-J, Bickert AA, Brown SP. 2019. The Synaptic Organization of Layer 6 Circuits Reveals Inhibition as a Major Output of a Neocortical Sublamina. Cell Rep. 28:3131-3143.e5.

Gilbert CD, Wiesel TN. 1979. Morphology and intracortical projections of functionally characterised neurones in the cat visual cortex. Nature. 280:120–125.

Guo W, Clause AR, Barth-Maron A, Polley DB. 2017. A Corticothalamic Circuit for Dynamic Switching between Feature Detection and Discrimination. Neuron. 95:180-194.e5.

Hains BC, Saab CY, Waxman SG. 2006. Alterations in burst firing of thalamic VPL neurons and reversal by Na(v)1.3 antisense after spinal cord injury. J Neurophysiol. 95:3343– 3352.

Halassa MM, Chen Z, Wimmer RD, Brunetti PM, Zhao S, Zikopoulos B, Wang F, Brown EN, Wilson MA. 2014. State-dependent architecture of thalamic reticular subnetworks. Cell. 158:808–821.

Han X, Qian X, Bernstein JG, Zhou H-H, Franzesi GT, Stern P, Bronson RT, Graybiel AM, Desimone R, Boyden ES. 2009. Millisecond-timescale optical control of neural dynamics in the nonhuman primate brain. Neuron. 62:191–198.

Hangya B, Ranade SP, Lorenc M, Kepecs A. 2015. Central cholinergic neurons are rapidly recruited by reinforcement feedback. Cell. 162:1155–1168.

Jurgens CWD, Bell KA, McQuiston AR, Guido W. 2012. Optogenetic stimulation of the corticothalamic pathway affects relay cells and GABAergic neurons differently in the mouse visual thalamus. PLoS One. 7:e45717.

Kim J, Matney CJ, Blankenship A, Hestrin S, Brown SP. 2014. Layer 6 corticothalamic neurons activate a cortical output layer, layer 5a. J Neurosci. 34:9656–9664.

Kirchgessner MA, Franklin AD, Callaway EM. 2020. Context-dependent and dynamic functional influence of corticothalamic pathways to first- and higher-order visual thalamus. Proc Natl Acad Sci U S A. 117:13066–13077.

Kvitsiani D, Ranade S, Hangya B, Taniguchi H, Huang JZ, Kepecs A. 2013. Distinct behavioural and network correlates of two interneuron types in prefrontal cortex. Nature. 498:363–366.

Lam Y-W, Sherman SM. 2010. Functional organization of the somatosensory cortical layer 6 feedback to the thalamus. Cereb Cortex. 20:13–24.

Lee CC, Sherman SM. 2008. Synaptic properties of thalamic and intracortical inputs to layer 4 of the first- and higher-order cortical areas in the auditory and somatosensory systems. J Neurophysiol. 100:317–326.

Lee CC, Sherman SM. 2009. Glutamatergic inhibition in sensory neocortex. Cereb Cortex. 19:2281–2289.

Lee CC, Sherman SM. 2012. Intrinsic modulators of auditory thalamocortical transmission. Hear Res. 287:43–50.

Madisen L, Mao T, Koch H, Zhuo J-M, Berenyi A, Fujisawa S, Hsu Y-WA, Garcia AJ, Gu X, Zanella S, Kidney J, Gu H, Mao Y, Hooks BM, Boyden ES, Buzsáki G, Ramirez JM, Jones AR, Svoboda K, Han X, Turner EE, Zeng H. 2012. A toolbox of Cre-dependent optogenetic transgenic mice for light-induced activation and silencing. Nat Neurosci. 15:793–802.

Mease RA, Krieger P, Groh A. 2014. Cortical control of adaptation and sensory relay mode in the thalamus. Proc Natl Acad Sci U S A. 111:6798–6803.

Olsen SR, Bortone DS, Adesnik H, Scanziani M. 2012. Gain control by layer six in cortical circuits of vision. Nature. 483:47–52.

Onodera K, Kato HK. 2022. Translaminar recurrence from layer 5 suppresses superficial cortical layers. Nat Commun. 13:1–16.

Payeur A, Guerguiev J, Zenke F, Richards BA, Naud R. 2021. Burst-dependent synaptic plasticity can coordinate learning in hierarchical circuits. Nat Neurosci. 1–10.

Potez S, Larkum ME. 2008. Effect of common anesthetics on dendritic properties in layer 5 neocortical pyramidal neurons. J Neurophysiol. 99:1394–1407.

Sceniak MP, Maciver MB. 2006. Cellular actions of urethane on rat visual cortical neurons in vitro. J Neurophysiol. 95:3865–3874.

Schmitt LI, Wimmer RD, Nakajima M, Happ M, Mofakham S, Halassa MM. 2017. Thalamic amplification of cortical connectivity sustains attentional control. Nature. 545:219– 223.

Sherman SM, Guillery RW. 2002. The role of the thalamus in the flow of information to the cortex. Philos Trans R Soc Lond B Biol Sci. 357:1695–1708.

Whilden CM, Chevée M, An SY, Brown SP. 2021. The synaptic inputs and thalamic projections of two classes of layer 6 corticothalamic neurons in primary somatosensory cortex of the mouse. J Comp Neurol.

Yu J, Hu H, Agmon A, Svoboda K. 2019. Recruitment of GABAergic Interneurons in the Barrel Cortex during Active Tactile Behavior. Neuron.

Zhan XJ, Cox CL, Rinzel J, Sherman SM. 1999. Current clamp and modeling studies of low-threshold calcium spikes in cells of the cat’s lateral geniculate nucleus. J Neurophysiol. 81:2360–2373.

Ziegler K, Folkard R, Gonzalez AJ, Burghardt J, Antharvedi-Goda S, Martin-Cortecero J, Isaías-Camacho E, Kaushalya S, Tan LL, Kuner T, Acuna C, Kuner R, Mease RA, Groh A. 2023. Primary somatosensory cortex bidirectionally modulates sensory gain and nociceptive behavior in a layer-specific manner. Nat Commun. 14:2999.

